# Modeling of Intracellular Taurine Levels Associated with Ovarian Cancer Reveals Activation of p53, ERK, mTOR and DNA-damage-sensing-dependent Cell Protection

**DOI:** 10.1101/2023.02.24.529893

**Authors:** Daniel Centeno, Sadaf Farsinejad, Elena Kochetkova, Tatiana Volpari, Alexandra Gladych-Macioszek, Agnieszka Klupczynska – Gabryszak, Teagan Polotaye, Michael Greenberg, Douglas Kung, Emily Hyde, Sarah Alshehri, Tonja Pavlovic, William Sullivan, Szymon Plewa, Helin Vakifahmetoglu-Norberg, Frederick J Monsma, Patricia A. J. Muller, Jan Matysiak, Mikolaj Zaborowski, Analisa DiFeo, Erik Norberg, Laura A. Martin, Marcin Iwanicki

## Abstract

Taurine, a non-proteogenic amino acid, and commonly used nutritional supplement can protect various tissues from degeneration associated with the action of the DNA-damaging chemotherapeutic agent cisplatin. Whether and how taurine protects human ovarian cancer (OC) cells from DNA damage caused by cisplatin is not well understood. We have found that OC ascites-derived cells contained significantly more intracellular taurine than cell cultures modeling OC. In culture, elevation of intracellular taurine concentration to OC ascites-cells-associated levels suppressed proliferation of various OC cell lines and patient-derived organoids, reduced glycolysis, and induced cell protection from cisplatin. Taurine cell protection was associated with decreased DNA damage in response to cisplatin. A combination of RNA sequencing, reverse phase protein arrays, live-cell microscopy, flow cytometry, and biochemical validation experiments provided evidence for taurine-mediated induction of mutant- or wild-type p53 binding to DNA, and activation of p53 effectors involved in negative regulation of the cell cycle (p21), and glycolysis (TIGAR). Paradoxically, taurine’s suppression of cell proliferation was associated with activation of pro-mitogenic signal transduction including ERK, mTOR, and increased mRNA expression of major DNA damage sensing molecules such as DNAPK, ATM and ATR. While inhibition of ERK or p53 did not interfere with taurine’s ability to protect cells from cisplatin, suppression of mTOR with Torin2, a clinically relevant inhibitor that also targets DNAPK and ATM/ATR, broke taurine’s cell protection. Our studies implicate that elevation of intracellular taurine could suppress cell growth, metabolism, and activate cell protective mechanisms involving mTOR and DNA damage sensing signal transduction.

## INTRODUCTION

Taurine is a non-proteogenic sulfonic amino acid implicated in tissue regeneration [1]. Taurine’s levels drop significantly with aging [2], and human clinical studies have demonstrated that taurine supplementation can treat congestive heart failure [3], retinal degeneration, and cardiomyopathy [1]. Studies using various animal models showed that taurine improved cognitive functions [4], glucose metabolism [5], motor functions [6], and protected various tissues from degeneration associated with cisplatin treatment [7, 8]. At the molecular level, taurine’s cell protection has been linked to decreased DNA damage, and chromatin remodeling [2]. These studies established a possible role for taurine in managing age-related or treatment-induced morbidities linked to DNA damage.

The tumor suppressor p53 protein, a product of the *TP53* gene, is a transcription factor that activates a plethora of signal transduction mechanisms involved in cell-cycle arrest, regulation of metabolism, DNA damage resolution, and cell survival under stress [9–11]. During DNA damage, p53 is rapidly stabilized through the ATM/ATR-dependent negative regulation of E3 ubiquitin ligase MDM2 [12], and the phosphorylation by DNAPK [13]. Stabilized p53 can suppress the cell cycle, trigger DNA repair, and rewire metabolism to support cell regeneration [14, 15]. *TP53* mutations are common in OC [16], and studies linked loss-of-p53 functions with increased carcinoma cell proliferation and cancer progression [17] . Paradoxically, however, in humans the prevalence of *TP53* mutations among non-malignant epithelial cells can be high [18], suggesting the existence of factors that support p53-growth-suppressive activities despite the mutations.

Target of rapamycin (TOR) including its mammalian ortholog mTOR is a serine/threonine kinase that is essential for cell proliferation, cell survival, and autophagy [19]. mTOR phosphorylation of ribosomal-6-kinase leads to activation of ribosomal protein S6 and translation initiation factor eIF4B [20], which ultimately regulates protein translation [21]. mTOR can be activated by signal transduction pathways downstream of Ras [22], and chemical biology approaches have been used to target mTOR in cancers [23]. For instance, inhibition of mTOR with the rapamycin derivative everolimus increases cisplatin sensitivity of cancer cells upon p53 pathway restoration [24], implicating that a combination of mTOR inhibition and reactivation of wild type functions in mutant p53 proteins might provide an additional strategy to increase sensitivity of cancer cells to standard therapy, such as cisplatin.

In this study, we report levels of intracellular pools of the free amino acid taurine in ascites cells derived from OC patients not previously administered chemotherapeutic treatment. Mimicking intracellular taurine levels in cultured cell lines and supplementing ovarian patient-derived organoids with taurine revealed taurine-mediated increase in cell size, inhibition of proliferation, suppression of glycolysis, and cell protection from DNA damage agent cisplatin. Combined RNA-sequencing, reverse-phase protein array (RPPA), western blotting and flow cytometry analysis of taurine-supplemented cell cultures underscored p53, ERK, mTOR/DNAPK/ATM/ATR pathway activation, and suppression of cisplatin-induced DNA damage as measured by γH2AX, and DNAPK phosphorylation. Inhibition of mTOR/DNAPK/ATM/ATR (Torin2), but not MEK (PD03255901), S6K1 (PF-4708671) or p53 expression (shRNA) induced death of taurine supplemented cultures in the presence of cisplatin. Our data support the idea that elevation of intracellular free-amino acid taurine can activate growth suppressive and cell protective signal transduction pathways involving p53, and mTOR/DNAPK/ATM/ATR.

## MATERIALS and METHODS

### Cell culture

FNE cells (kind gift from Dr. Tan Ince, Weill Cornell Medicine, New York) were cultured in a 1:1 ratio of DMEM/F12 (HiMedia), and Medium 199 (HiMedia), 2% heat-inactivated Fetal Bovine Serum (HI-FBS; Sigma-Aldrich), 1% *v/v* penicillin-Streptomycin (VWR), 0.5 ng/mL of beta-estradiol (US Biological), 0.2 pg/mL of triiodothyronine (Sigma-Aldrich), 0.025 μg/mL all-trans retinoic acid (Beantown Chemical), 14 μg/mL of insulin (Sigma-Aldrich), 0.5 ng/mL of EGF (Peprotech), 0.5 μg/mL hydrocortisone (Sigma-Aldrich), and 25 ng/mL of cholera toxin (Calbiochem). Kuramochi, OVCAR4, and OV90 (kind gift from Dr. Denise Connolly, Fox Chase Cancer Center) and HEK293T cells (Thermo Fisher) were cultured in DMEM/F12 supplemented with 10% HI-FBS and 1% v/v penicillin-Streptomycin. CaOv3 (ATCC), Hey-A8 (kind gift from Dr. Sumegha Mitra’s laboratory, University of Indiana), and TYK-nu (kind gift from Dr. Joan Brugge’s laboratory, Harvard Medical School) cells were cultured in a 1:1 ratio of MCDB 105 (Sigma-Aldrich) and Medium 199 supplemented with 5% HI-FBS and 1% v/v penicillin-Streptomycin. OV81-CP, OV81-CP40 and OV231, OV231-CP30 were cultured in DMEM/F12 supplemented with 10% HI-FBS and 1% v/v penicillin-Streptomycin. All cell lines were cultured in a humidified incubator at 37°C and with 5% carbon dioxide. Cell cultures were tested for the presence of mycoplasma every 3-6 months using the Uphoff and Drexler detection method [25]. Taurine (TCI Chemicals) was dissolved directly into complete cell culture media at a concentration of 40 mg/mL and passed through a 0.22 micron filter before treatment.

### Patient Ascites

For a study from 2021 to 2022, we enrolled Caucasian patients with histologically confirmed high-grade serous OC and ascites in the Department of Gynecologic Oncology, Poznań University of Medical Sciences. Ovarian tumors were staged according to FIGO (International Federation of Gynecology and Obstetrics) system. Ascites fluid samples were collected from patients (i) at laparoscopy before starting neoadjuvant chemotherapy. Ascites fluid (10 mL) was centrifuged within 2 hours after collection in a falcon tube at 1,100 × *g* for 10 minutes at room temperature to separate a cell pellet. The supernatant and malignant cell pellets were stored in -80°C and analyzed collectively. All patients provided a signed informed consent, approved by the Ethics Review Board of Poznań University of Medical Sciences (Consent No 737/17). All procedures performed in studies involving human participants were in accordance with the ethical standards of the institutional and national research committee and with the 1964 Helsinki declaration and its later amendments or comparable ethical standards.

### Mass-spectrometry-based quantification of taurine levels in ascites and cell lysates

The panel of amino acids was quantified based on aTRAQ kit for amino acid analysis (SCIEX, Framingham, MA, USA) and liquid chromatography coupled to a triple quadrupole tandem mass spectrometry technique. The samples (ascites or cell lysates) were thawed at room temperature, and 40 μL of a matrix was transferred to a 0.5 mL Eppendorf tube. Then, 10 μL of sulfosalicylic acid was added to precipitate the proteins, and the vial contents were mixed and centrifuged. Subsequently, 10 μL of supernatant was transferred to a clean tube, and 40 μL of borate buffer was added, mixed, and centrifuged. In the next step, the 10 μL of the obtained mixture was transferred to a clean tube and mixed with 5 μL of amino-labeling reagent (aTRAQ Reagent Δ8®). After centrifugation, samples were incubated for 30 minutes at room temperature. The incubation was followed by the addition of 5 μL of hydroxylamine solution, mixing and centrifugation. Then the samples were incubated for 15 minutes at room temperature. In the next step, 32 μL of freshly prepared internal standards solution was added, mixed up, and centrifuged. The contents of the tubes were concentrated (temperature 50°C for about 15 minutes) to a volume of about 20 μL using a vacuum concentrator (miVac Duo, Genevac, Stone Ridge, NY, USA). In the last step, 20 μL of ultrapure water was added to each vial and mixed. The contents of the tubes were transferred to amber-glass autosampler vials with inserts. Samples were analyzed in random order by chromatographic separation followed by tandem mass spectrometry detection LC-MS/MS. The analytes were separated on a Sciex C18 column (4.6 mm × 150 mm, 5 μm) maintained at 50°C using a 1260 Infinity HPLC instrument (Agilent Technologies, Santa Clara, CA, USA). A gradient flow of the mobile phase was applied. The mobile phase consisted of 0.1% formic acid (FA) and 0.01% heptafluorobutyric acid (HFBA) in water—phase A, and 0.1% FA and 0.01% HFBA in methanol—phase B, maintained at a flow rate 800 μL/min. Total runtime was 18 minutes per sample, with injection volume equal to 2 μL. Detection and quantitation of analytes were performed by means of a quadrupole tandem mass spectrometer with an electrospray ionization (ESI) TurboV ion source operated in positive-ion mode. All results were generated in a scheduled multiple reaction monitoring mode. Raw data from amino-acid assays was acquired and analyzed using the Analyst software version 1.6.3 (Sciex, Framingham, MA, USA). The method validation and sample preparation were described in detail before [26].

### Quantification of taurine levels in cell lysates using a commercial kit

To determine intracellular taurine concentrations, a Taurine Assay Kit (Cell Biolabs, Inc.) was used. Cells were treated with taurine for the indicated amount of time, washed three times with PBS, lysed in ice-cold RIPA buffer to extract intracellular contents, and protein content was determined using BCA Assay. Then, 25 μg of protein were loaded into a 96-well plate in triplicate for each sample. Taurine concentration was determined using the Taurine Assay Kit following the manufacturer’s instructions, and absorbance values were read at 405 nm using a SpectraMax i3x Multi-Mode Microplate Reader (Molecular Devices). Absorbance values were then background subtracted and normalized to the average absorbance value of the control group. Values were reported as fold change in intracellular taurine concentration.

### Organoid derivation and culture

Organoids were derived from primary tissue tumor samples of patients with OC as described previously [27]. Briefly, fresh tumor resections were cut in small pieces. Two to four random pieces were snap frozen and stored in liquid nitrogen for molecular analysis or fixed for histology. The remaining tissue was minced, digested with 0.7 mg/mL collagenase (Sigma C9407) in the presence of 10 mM ROCK inhibitor (Y-27632 dihydrochloride, Abmole) at 37°C for 25-50 minutes and strained over a 100-μm filter. The digested tissue suspension was centrifuged at 300 × *g* for 5 minutes. Visible red pellets were lysed with red-blood lysis buffer (Sigma-Aldrich) for 3 minutes at room temperature and centrifuged at 300 x *g* for 5 minutes. The dissociated tissue pellet was resuspended in cold Cultrex Reduced Growth Factor BME type 2 (Cultrex BME) (R&D systems) and 25 mL drops per well were plated on pre-warmed 48-well plates. The Cultrex BME/cell suspension droplets were allowed to solidify at 37°C for at least 30 minutes. Solidified droplets with the embedded cells were then overlaid with ovarian tumor organoid medium containing 10 mM ROCK inhibitor (Y-27632 dihydrochloride, Abmole). Medium is composed of: Advanced DMEM/F12 (Gibco) containing GlutaMAX (Gibco), 100 U/mL penicillin, 100 mg/mL streptomycin (Gibco), 10 mM HEPES (Gibco), 100 mg/mL Primocin® (InvivoGen), 1x B-27™ supplement (Gibco), 1.25 mM N-Acetyl-L-cysteine (Sigma-Aldrich), 10 mM Nicotinamide (Sigma-Aldrich), 0.5 mM A83-01 (R&D systems), 0.5 mg/mL Hydrocortisone (Sigma-Aldrich), 10 mM Forskolin (R&D systems), 100 nM b-Estradiol (Sigma-Aldrich), 16.3 mg/mL BPE (Thermo Fisher Scientific), 10 ng/mL recombinant human FGF-10 (PeproTech), 5 ng/mL recombinant human FGF-7 (PeproTech), 37.5 ng/mL recombinant human Heregulin Beta-1 (PeproTech), 5 ng/mL recombinant human EGF (PeproTech), 0.5 nM WNT Surrogate-Fc fusion protein (ImmunoPrecise), 10% R-Spondin1 conditioned medium and 1% Noggin-Fc fusion protein conditioned medium (ImmunoPrecise). Media was changed every 2-3 days. For established lines, dense cultures containing organoids ranging in size from 100 to 500 µm were passaged every 7-10 days with TrypLE (Life Technologies) containing 10 mM ROCK inhibitor and 10 mg/mL DNase I (Sigma-Aldrich). For taurine supplementation, 2- aminoethanesulfonic acid (TCI chemicals) was freshly added directly to ovarian tumor organoid medium at a 160 mM final concentration, filtered and stored at 4°C. Filtered ovarian tumor organoid medium from the same batch was used as control. For single-cell plating, dense organoid cultures were dissociated with TrypLE (Life Technologies) containing 10 mM ROCK inhibitor and 10 mg/mL DNase I (Sigma-Aldrich) for 10 minutes at 37°C followed by mechanical shearing through a 30-G needle, 5x dilution with Advanced DMEM/F12 (Gibco) containing antibiotics, HEPES and GlutaMax, as indicated above supplemented with 10 mM ROCK inhibitor and 10 mg/mL DNase I, and centrifugation at 300 × *g* for 5 minutes. Cell pellets were resuspended in cold Cultrex BME and plated at 400 cells/mL of BME. Media ± taurine was fully changed every 2-3 days. For organoid collection and plating, 20-50 µm organoids were extracted from the Cultrex BME droplet using cold Cell Recovery Solution containing 10 mg/mL DNase I at 4°C for 30 minutes. The dissolved Cultrex BME containing intact organoids was then collected into 2-3x volume of cold Advanced DMEM/F12 containing 0.1% BSA (Sigma-Aldrich) and 10 mg/mL DNase I and centrifuged at 200-250 × *g* for 5 minutes. The organoid solution is resuspended in cold ovarian tumor organoid medium containing 10% Cultrex BME with or without taurine. The organoid solution is counted prior plating at 500 organoids/well in 384-well plate format.

### Organoid metrics

Bright-field images were acquired and analyzed by a Perkin Elmer Opera Phenix High-Content Screening System and Harmony high-content analysis software (4.9). The area of each organoid was measured with an automated quantification pipeline set up to identify well defined organoid structures in the same focal plane of one single z-plane after selection based on image texture, single structure size, and morphology.

### Organoid ATP assay

CellTiter-Glo® 3D Cell Viability Assay (Promega) was performed on Patient-Derived Organoids (PDOs) to measure ATP in metabolically active cells at the specified experimental endpoint following the manufacturer’s instructions. Briefly, CellTiter-Glo® 3D Reagent was added to each well at 1:1 ratio. Plates were shaken for 25 minutes and luminescence was measured with a CLARIOstar® Plus reader (BMG LABTECH).

### Cell proliferation analysis

Cell proliferation was determined using an automated cell counter or by live cell imaging. To assess proliferation using an automated cell counter, cells were first seeded into a 12-well plate. The following day, three wells per condition were treated as indicated and another three wells were trypsinized and counted using a Luna II Automated Cell Counter (Logos Biosystems). After 72h of treatment, cells were once again harvested and counted, and cell numbers were normalized to the average value of the initial count and reported as fold change in cell number.

To determine proliferation by live cell imaging, GFP-expressing FNE-m-p53 cells were seeded in a 6-well plate and captured over time using a BioTek Lionheart FX Automated Microscope (BTLFX). Images were analyzed using the ImageJ TrackMate plugin.

### Cell volume and adhesion area quantification

GFP-expressing FNE-m-p53 cells were seeded in a 6-well plate and incubated overnight. The following day, cells were treated as indicated and incubated for 72h. Cells were then trypsinized and seeded into a low-adhesion 96-well round bottom plate at a density of 100 cells per well. The plate was then centrifuged at 900 rpm for 5 minutes and cell clusters were immediately imaged using a BTLFX. Images were captured using a 10X objective in the GFP- and phase-contrast channels. To determine cell volume, GFP images were processed in FIJI using an in-house macro. Briefly, images were background subtracted and an automated threshold was applied. Images were then converted to binary and a watershed was applied to separate adjacent cells. The area of each particle (cell) was then measured. From this value, the radius of each cell was determined and volume was estimated from the equation for the volume of a sphere (V = 4/3 π r³). Between 15-20 wells were analyzed per condition, and the average volume of a cell per well was reported.

To quantify cell adhesion area, cells were seeded and treated as described for the proliferation assay. At the endpoint, images were captured using a BTLFX with a 10X objective in the GFP- and phase-contrast channels. Then, as described for the fluorescence microscopy cell proliferation assay, images were processed in FIJI with an in-house macro and the area of each particle (cell) was reported as cell adhesion area.

### Basement-membrane reconstitution and spheroid size quantification

To examine the effects of taurine on basement-membrane (BM) reconstituted organotypic structures, FNE-m-p53, Hey-A8, and OV90 cells were reconstituted in Matrigel® Growth Factor Reduced Basement Membrane Matrix (MG) (Corning) as previously described [28]. One hundred cells were seeded per well in a 96-well ultra-low adhesion Nunclon Sphera™ plate, centrifuged at 900 rpm for 5 minutes, and incubated overnight. The following day, on ice, MG was added to cell culture media to a concentration of 4% v/v. MG solution was then added to each well containing 100 μL of clustered spheroids to a final volume of 200 μL and MG concentration of 2% v/v. Z-projections of organotypic structures were captured at the indicated time points using a BTLFX using the 4X objective in the bright-field and GFP channels. Z-projections were processed into focused stacks using Gen5 software and the area of each structure was measured using FIJI.

### RNA Isolation and targeted RT-qPCR analysis

Total RNA was isolated using Quick-RNA Miniprep Kit (Zymo) and reverse transcribed into cDNA using High Capacity cDNA Reverse Transcription Kit (Applied Biosystems) following the manufacturer’s instruction. To analyze mRNA expression, 10 ng of cDNA was added to a 10 μL reaction mixture containing Power SYBR Green Master Mix (Applied Biosystems) and primers at a concentration of 330 nM designed to detect the transcript of interest. mRNA levels were quantified using a CFX96 Touch Deep Well Real-Time PCR Detection System (Bio-Rad) and mRNA expression was normalized to a housekeeping gene.

### RNA sequencing and transcriptomic analysis

RNA extraction was performed by E.Z.N.A.® total RNA kit (Omega Bio-tek) following the manufacturer’s protocol. RNA concentration and purity was assessed by NanoDrop 2000 (ThermoFisher). RNA samples were stored at -80°C freezer and subsequently shipped on dry ice to Genewiz (Azenta US, South Plainfield, New Jersey) for RNA sequencing. The RNA sequencing library preparation workflow started with PolyA–based mRNA enrichment, mRNA fragmentation, and random priming followed by first- and second-strand complementary DNA (cDNA) synthesis. Subsequently, end-repair with 5′ phosphorylation and adenine (dA)–tailing was carried out. Finally, adaptor ligation, PCR enrichment, and Illumina HiSeq 2500 sequencing with two 150–base-pair (bp) runs were performed, and sequence reads were mapped to the reference genome. Bioinformatics analysis workflow started with evaluation of sequence quality and trimming the reads by means of Trimmomatic v.0.36 software to remove possible adapter sequences and nucleotides with poor quality. STAR aligner v.2.5.2b software was used to map the trimmed reads to the *Homo sapiens* GRCh38 reference genome available on ENSEMBL. To extract gene hit counts, feature counts from the Subread package v.1.5.2. were used. The hit counts were reported based on the gene ID feature in the annotation file. Only reads that were within the exon regions were considered. The table generated from extracted gene hit counts was used for differential expression analysis. Using DESeq2, gene-expression comparison between the taurine-treated and control samples was performed. By using the Wald test, p-values and log2-fold changes were calculated. Genes with an adjusted p-value (Padj) of less than 0.05 and absolute log2-fold change more than 1 were counted as differentially expressed genes (DEG).

Gene ontology analysis was applied on the statistically significant genes with GeneSCF v.1.1-p2 software. The goa_human GO list was referenced to cluster the set of genes based on their biological functions and determine their statistical significance. Gene set enrichment analysis (GSEA) was performed, as described earlier (Subramanian et al., 2005), with GSEA_4.2.2. software, against selected gene sets from Molecular Signatures Database (MSigDB 7.5). GSEA was performed with the ranking metric set to Signal2noise and with the number of permutations set to 1000. Plots with false discover rate (FDR) of less than 0.25 were considered significant.

### Flow cytometry

DNA staining (cell-cycle analysis) by flow cytometry was performed as follows. Cells were harvested by trypsinization, collected by centrifugation, and washed twice with PBS. Cell pellets were then fixed with ice-cold 70% ethanol for at least 30 minutes on ice. Cells were then washed twice with PBS and treated with PBS containing 100 μg/mL RNAse A (Sigma-Aldrich) and 2 μg/mL propidium iodide (PI) (Molecular Probes) for 30 minutes at room temperature in the dark before analysis. Data acquisition was performed using an Attune NxT Flow Cytometer (ThermoFisher). Data were then analyzed using FlowJo software.

To determine cell viability, cell culture supernatant was collected, adherent cells were trypsinized, and were centrifuged at 300 × g for 5 minutes. Cells were washed twice with PBS containing 2% FBS. Cells were then incubated on ice with PBS containing 2% FBS and 2 μg/mL PI. Data was then acquired by flow cytometry, analyzed using FlowJo, and viability was reported as the percentage of PI-positive cells.

To stain for phosphorylated γH2AX, cells were treated for 48 hours and collected by trypsinization. Cells were washed and fixed in 4% paraformaldehyde (PFA) for 15 minutes, followed by permeabilization in ice-cold methanol for 30 minutes. Cells were stained with Phospho-Histone H2A.X primary antibody (Cell Signaling Technology #2577) at dilution of 1:200 in 0.5% BSA dissolved ion PBS for one hour at room temperature. Cells were then washed and stained with Alexa Fluor 568 secondary antibody (Invitrogen A-11011) at a dilution of 1:500. Cells were then analyzed by flow cytometry.

### Plasmids

SLC6A6 shRNA plasmids were obtained from Dharmacon (pLKO.1 vector; TRCN0000038409: TATCACCTCCATATATCCAGG, referred to in this manuscript as G5, and TRCN0000038412: TATACTTGTACTTGTTGTAGC, referred to in this manuscript as G8).

To generate a p21-reporter plasmid, NEBuilder HiFi DNA Assembly was used to clone a p21 promoter sequence upstream of mCherry-3X-NLS into a lentiviral vector. A sequence containing the 2.4-kbp p21 promoter region was PCR amplified from the pGL2-p21 promoter-Luc (Addgene #33021). The mCherry-3X-NLS sequence was PCR amplified from CSII-prEF1a- mCherry-3X-NLS (Addgene #125262). PCR fragments were assembled into pLenti6.3/V5- DEST-GFP (Addgene #40125) by replacing the CMV promoter and EGFP sequences at the 5’ ClaI and 3’ SalI restriction sites, respectively. The resulting plasmid (pLenti6.3-p21 promoter-mCherry-3X-NLS) contains the 2.4-kbp p21 promoter upstream of the mCherry-3X-NLS sequence. Plasmid constructs were verified by Next Generation Sequencing.

### Lentivirus production

To generate lentivirus, packaging plasmids psPAX2 (Addgene #12260) and pMD2.G (Addgene #12259) were mixed with a lentiviral plasmid containing a gene of interest or shRNA and incubated with Lipofectamine 3000 (Invitrogen) in serum-free Opti-MEM (Gibco) to generate liposomes containing plasmid DNA. The mixture was then added to HEK293T cells and media was refreshed the following day. Cell culture supernatant containing viral particles was collected 48 hours and 72 hours after transfection. Cell lines were then transduced by adding viral particles to media containing polybrene (Santa Cruz Biotechnology) at a concentration of 10 μg/mL and selected with an appropriate antibiotic 24 to 48 hours after transduction to generate cells stably expressing transgene or shRNA.

### Western blot analysis

Cells were washed twice with PBS and lysed in ice-cold RIPA buffer (Cell Signaling Technology) supplemented with a Halt^TM^ Protease Inhibitor Cocktail (Thermo Fisher), and cells were collected using a cell scraper. Cell lysates were pelleted at 20,000 x *g* for 20 minutes, supernatant was collected, and protein concentration was determined by BCA Assay (Pierce) according to the manufacturer’s instructions. Lysates were then mixed with 6X sample buffer (375 mM Tris Base, 9% sodium dodecyl sulfate, 50% glycerol, 0.075% bromophenol blue, 9% β-mercaptoethanol), boiled for 10 minutes at 95°C, and then loaded on a polyacrylamide gel and resolved by electrophoresis. Proteins were then transferred to Immobilon membranes (Whatman), blocked with 5% non-fat milk in Tris-buffered saline containing 0.1% *v/v* Tween-20 (TBST) for 30 minutes at room temperature. Membranes were incubated overnight at 4°C in primary antibodies diluted in 5% non-fat milk in TBST. Membranes were then washed three times with TBST and incubated in an HRP-conjugated secondary antibody (1:10,000) for 1 hour at room temperature. Membranes were then washed three times in TBST. Membranes were developed using Immobilon^TM^ Forte enhanced chemiluminescent substrate (Millipore) and visualized using an iBright CL1500 (Thermo Fisher).

### Reverse-phase protein array

Adherent FNE-m-p53 cells were cultured in triplicate for 72 hours in control media or with media supplemented with taurine. Cells were then washed three times on ice, and then lysed in RPPA lysis buffer (1% Triton-X-100, 50 mM HEPES, pH 7.4, 150 mM NaCl. 1.5 MgCl_2_ 1 mM EGTA, 100 mM NaF, 10 mM NaPPi, 10% glycerol, 1 mM Na_3_VO_4_, and Halt Protease Inhibitor (Thermo Fisher). Lysates were incubated on ice for 15 minutes, and then centrifuged at 13,000 x *g* for ten minutes. Following protein quantification, lysates were then diluted to equal concentration using RPPA buffer, and then boiled and denatured using 1% SDS for ten minutes. Lysates were then snap frozen and shipped on dry ice to MD Anderson Cancer Center for RPPA analysis.

### Analysis of glycolysis

Extracellular acidification rate (ECAR) was measured using an XFp Extracellular Flux Analyser (Seahorse Bioscience, North Billerica, MA, USA). CaOv3 cells were exposed to taurine for 48 hours and then reseeded (8000 cells/well) in taurine-containing media into the wells of an XFp Cell Miniplate. 24 hours later, the cells were analyzed using XFp Glycolysis Stress Test Kit (Seahorse Bioscience) following the manufacturer’s instructions.

Cells were washed 3 times with 100 µL of XF Base Media (Seahorse Bioscience) containing 2 mM L-Glutamine. Then the cells were incubated in 180 µL of the same media for 20 minutes. After three baseline ECAR measurements, cells were subsequently injected with 10LJmM glucose, 1 μM oligomycin, and 50LJmM 2-deoxyglucose (glycolysis inhibitor).

The data obtained were normalized to protein concentrations determined by the bicinchoninic acid (BCA) assay (Pierce, Thermo Fisher Scientific, Waltham, MA, USA). Wave (Agilent Technology) and GraphPad Prism 8 software were used to analyze and plot the data.

### Statistical analysis

Statistical analyses were performed using GraphPad Prism 9.0 (GraphPad Software, San Diego, CA, USA). Statistical significance was determined using an unpaired, two-tailed, parametric t-test, ordinary one-way ANOVA with Tukey’s *post hoc* multiple comparisons test, or two-way ANOVA with Sidak’s *post hoc* multiple comparisons test, with *p* ≤ 0.05 considered statistically significant.

## RESULTS

### Determination of intracellular taurine levels in patient-derived and tissue culture cells

To begin investigations of taurine’s cell-protective mechanisms, we first wanted to determine the taurine levels in cells derived from the ascites of OC patients. To do so, we utilized quantitative mass spectrometry to profile the intracellular levels of 36 free amino acids within OC ascites-derived cells obtained from six chemotherapy naïve patients (Fig. 1A). Our analysis established that taurine was one of the most abundant free amino acids among the selected intracellular pool of amino acids (Fig.1B). We found that intracellular taurine concentrations varied (45-900 μM) and averaged 430 μM.

**Figure 1.**
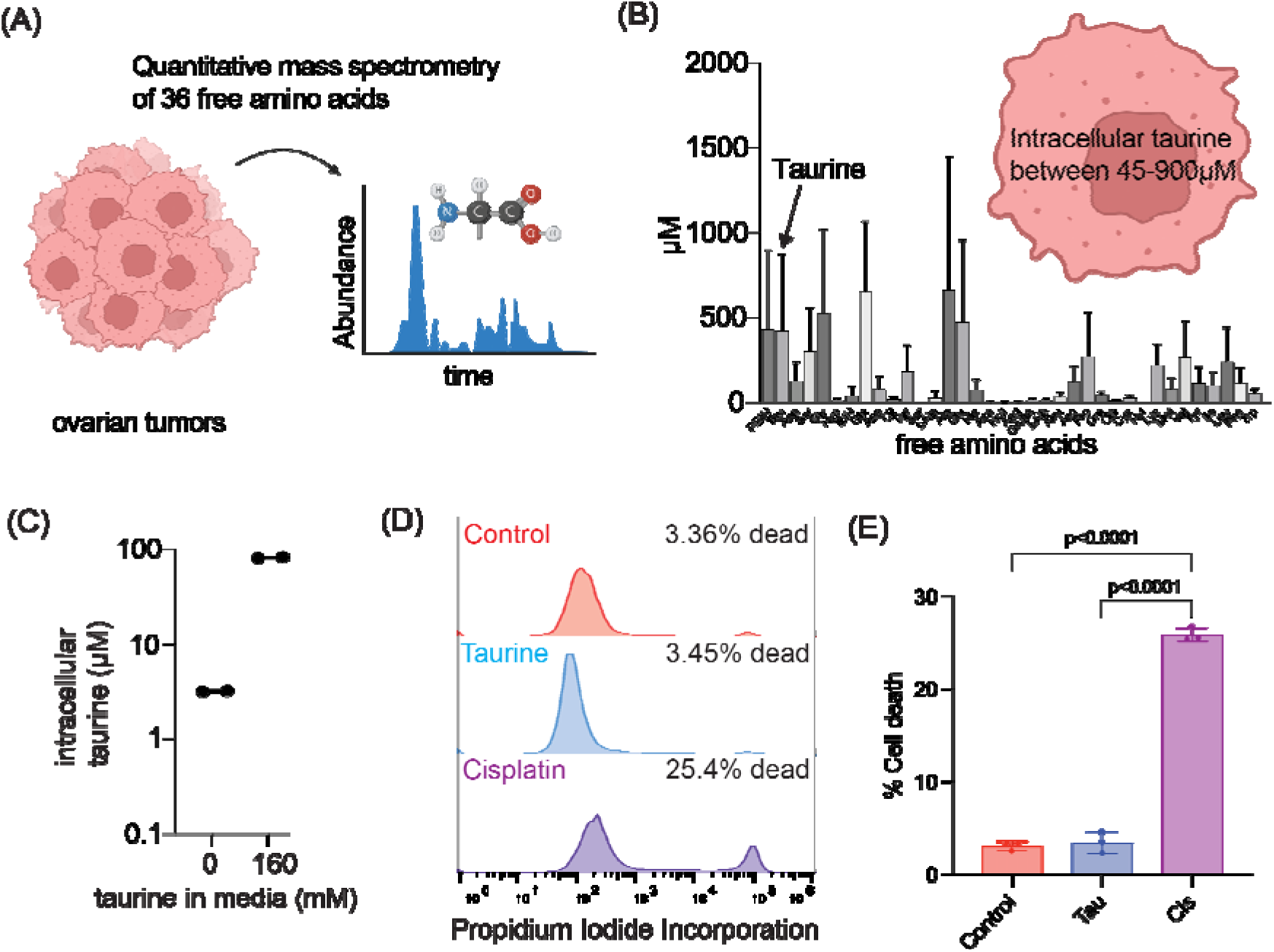
Modeling of intracellular taurine levels associated with OC in vitro. (A-B) Quantitative mass spectrometry workflow for the determination of free intracellular amino acid concentration in cells isolated from patient ascites. n=6 patient samples. Data are reported as mean ± SD. (C) Mass spectrometry quantification of intracellular taurine concentration in FNE-m-p53 cells cultured in control media or media supplemented with 160 mM taurine. Each dot is one replicate. (D) Representative histograms and (E) quantification of cell death for FNE- m-p53 cells treated with 160 mM taurine or 10 μM cisplatin for 72h. Viability was determined by propidium iodide (PI) staining and quantified using flow cytometry. n=3 replicates. Data are reported as mean ± SD and were

Since our tissue culture conditions do not include taurine supplementations, nor is taurine a component of commonly used OC cell culture media, we wanted to know the basal intracellular taurine levels in tissue culture cells. We began our experiments using fallopian tube non-ciliated epithelial (FNE) cells expressing OC driving mutant p53^R175H^ (m-p53) [29]. We chose this cell model for several reasons. First, FNE cells are considered to represent precursor cells for most of OCs [30]. Second, mutations in TP53 gene are nearly ubiquitous in OC disease [31]. Third, the taurine transporter SLC6A6 is negatively regulated by wild type p53 [32]. Fourth, as opposed to a majority of OC cell lines, FNE-m-p53 cells represent a chemotherapy naïve cell model, ruling out pre-existence of resistant mechanisms to DNA damaging agents. Using FNE- m-p53 as a cell culture model, we found that these cells contained, on average, 100 times less taurine compared to ascites-derived cells (Fig. 1C). Supplementation of FNE-m-p53 culture media with 160 mM taurine was sufficient to increase intracellular taurine content to within in the range of ascites cells (Fig. 1C) without causing cell death (Fig. 1D**-**E). Our data suggest that taurine is one of the most abundant intracellular amino acids in ascites cells isolated from OC patients, and that mimicking physiologic intracellular taurine concentration in cell culture is not toxic and can be accomplished by addition of taurine to media.

### shRNA-mediated attenuation of SLC6A6 expression decreases intracellular taurine levels in FNE-m-p53 cells

In most cells, taurine can be transported from outside *via* a sodium/chloride-dependent symporter, SLC6A6 [33]. So, we examined the functional consequences of SLC6A6 expression in the taurine transport of tissue culture cells. We first measured SLC6A6 mRNA expression using RT-qPCR in FNE-m-p53 cells expressing a control plasmid (pLKO vector) or plasmids containing SLC6A6 shRNA sequences (G5 and G8). We found that both shRNA plasmids depleted expression of SLC6A6 mRNA, with G8 being more efficient (Fig. 2A). We confirmed SLC6A6 shRNA G8 – mediated knock-down using a combination of cell membrane protein extraction and western blot analysis of FNE-m-p53 cell lysates (Fig. 2B). To examine the functionality of SLC6A6 knock-down, we used a commercially available taurine assay kit from Cell Biolabs (MET-5071) that measures intracellular taurine levels. We found that shRNA-mediated attenuation of SLC6A6 using the G8 shRNA induced a three-fold decrease in taurine intracellular levels (Fig. 2C). We verified these results using mass-spectrometry and found that, in FNE-m-p53 cells cultured in the presence of taurine, shRNA-mediated attenuation of SLC6A6 decreased intracellular taurine content from 130 μM to 48 μM (Fig. 2D). Our data support the idea that in cultured cells intracellular taurine accumulation is partially mediated by SLC6A6.

**Figure 2.**
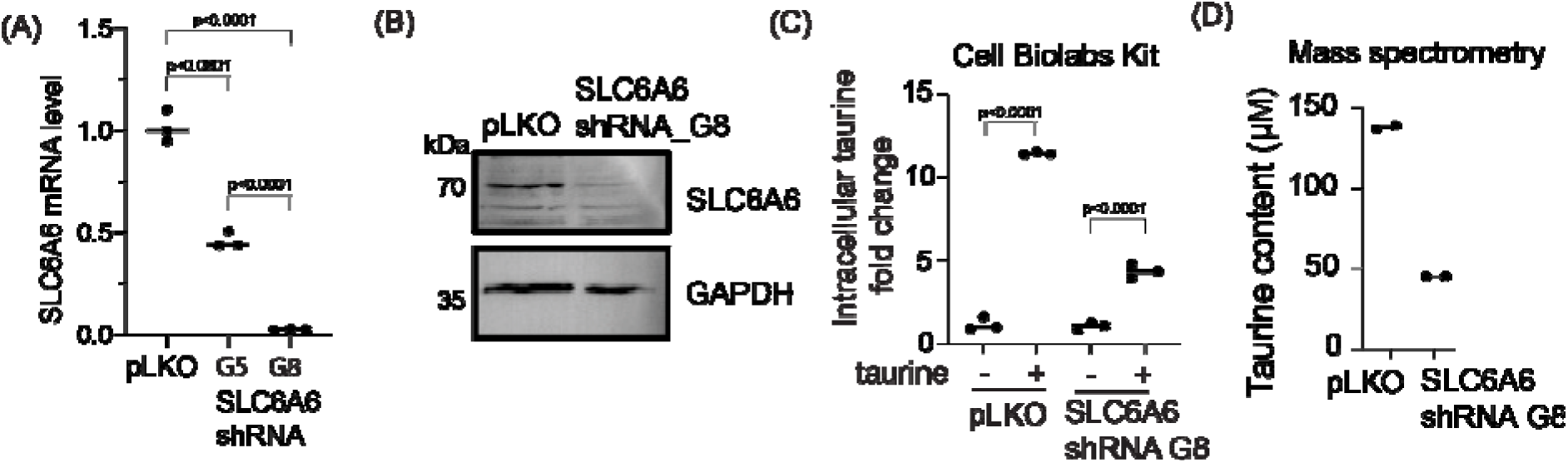
Intracellular taurine accumulation is mediated by the SLC6A6 transporter. (A) RT-q-PCR analysis of SLC6A6 mRNA expression in FNE-m-p53 cells transduced with scramble shRNA (pLKO) or SLC6A6 shRNA (G5 and G8). Each dot is one replicate. Statistical analysis was performed using one-way ANOVA. (B) Western blot of SLC6A6 expression in FNE-m-p53 cells transduced with scramble shRNA or SLC6A6 shRNA G8. (C) Fold change in intracellular taurine content following treatment with control media or media containing 160 mM taurine for 72h. Taurine concentration was determined using Cell Biolabs Taurine Assay Kit. Each dot is one replicate. Statistical analysis was performed using one-way ANOVA comparing no taurine *vs.* taurine groups. Line indicates median. (D) Mass spectrometry quantification of intracellular taurine content in cells treated with 160 mM taurine for 72h. Each dot is one replicate.

### Taurine supplementation inhibits OC cell proliferation in tissue culture

Our data suggest that exogenous taurine addition to culture media can elevate intracellular taurine in cultured cells to levels similar to cells in ascites without causing cell death (Fig. 1D-E). Based on these data we decided to propagate FNE-m-p53 and various OC cells in media containing 160 mM taurine, which can increase taurine intracellular levels to those observed in ascites-derived cells (Fig. 2D). Live-cell phase contrast micrography of taurine-supplemented FNE-m-p53 cell monolayer cultures showed fewer cells undergoing cell division and cell adhesion area increase (S. Fig. 1A-C, MOVIE 1). Because taurine is an osmolyte, and intracellular accumulation of osmolytes increases cell size and promotes G1 arrest [34], we hypothesized that elevation of intracellular taurine levels suppresses the proliferation of cultured cells. To test this hypothesis, we assessed the proliferation of multiple cell lines and cell culture models supplemented with taurine. We found that taurine supplementation suppressed cell proliferation in all tested cell lines, cultured as monolayers or, in some cases, under three- dimensional basement membrane reconstituted conditions (Fig. 3A-D). We also observed that growth suppression by taurine was associated with cell-cycle arrest (S. Fig. 1D-E), suggesting the activation of cell-cycle inhibitory programs.

**Figure 3.**
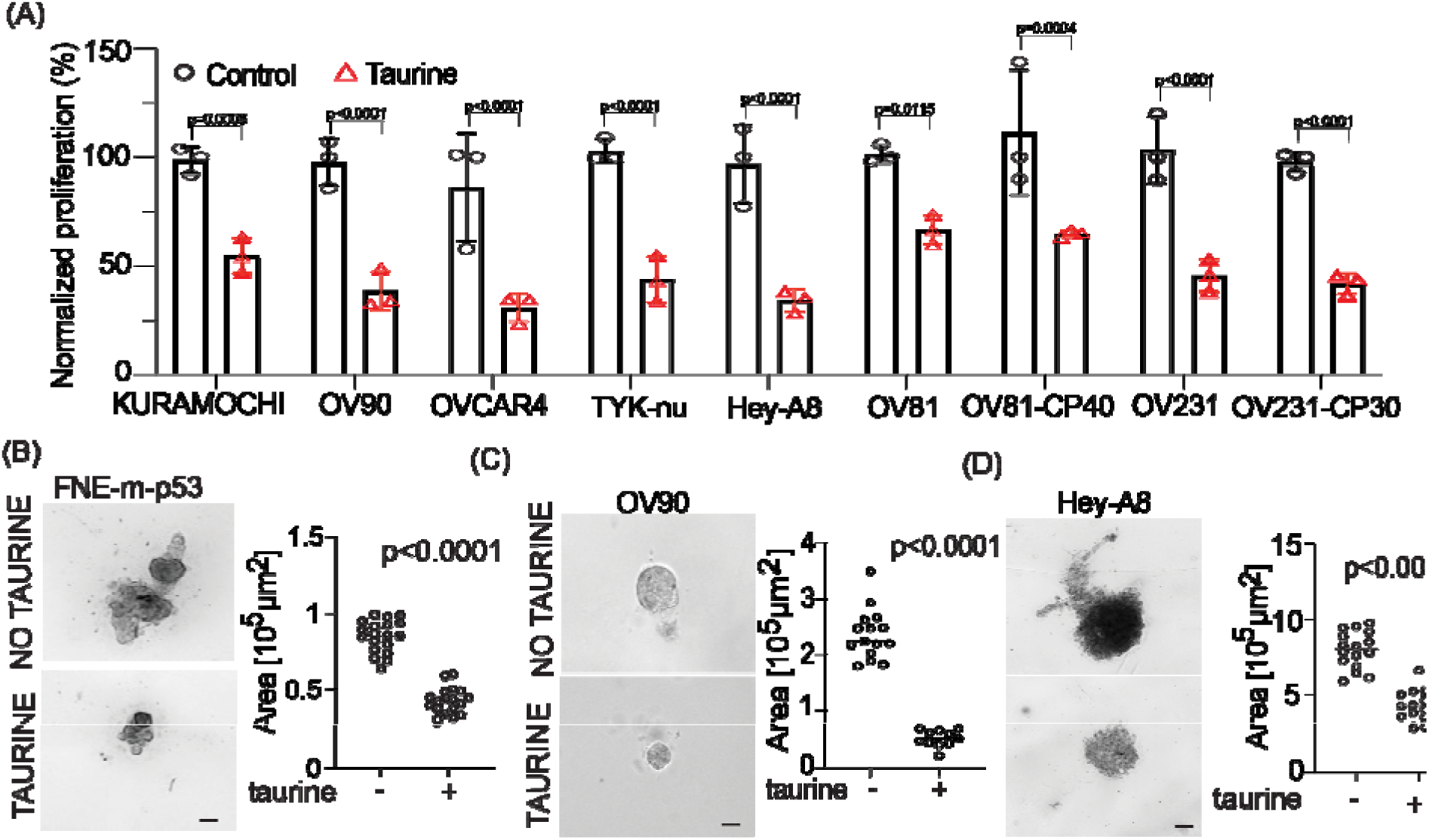
Taurine supplementation suppresses the growth of OC cell lines. (A) Proliferation of OC cells lines was quantified by cell counting. OC cells were treated with 160 mM taurine for 72h. Data was normalized to the mean cell count for each respective control group and is reported as mean ± SD. Each dot represents one replicate count. Statistical analysis was performed using Student’s t-test. (B-D) Bright-field microscopy images of basement membrane reconstituted organotypic structures. The size of each structure was quantified based on area. Each dot represents one structure, and the bar is the mean. Statistical analysis was determined using Student’s t-test. Scale bars are 50 µm.

Interestingly, taurine washout demonstrated phenotype reversal (S. Fig. 1F), supporting the idea that maintaining taurine pools is required for suppression of proliferation. Nevertheless, we were motivated by these experiments, and wanted to determine whether taurine supplementation suppresses the growth of patient-derived organoids (PDOs). We provide evidence that taurine significantly decreased the growth of various PDOs (Fig. 4A-B). Taurine’s PDO growth suppression was associated with a decrease in ATP production as measured by CellTiter-Glo® assays (Fig. 4C). Our data support the idea that taurine intracellular accumulation is associated with suppression of cell proliferation and the activation of cell-cycle and metabolic regulatory mechanisms.

**Figure 4.**
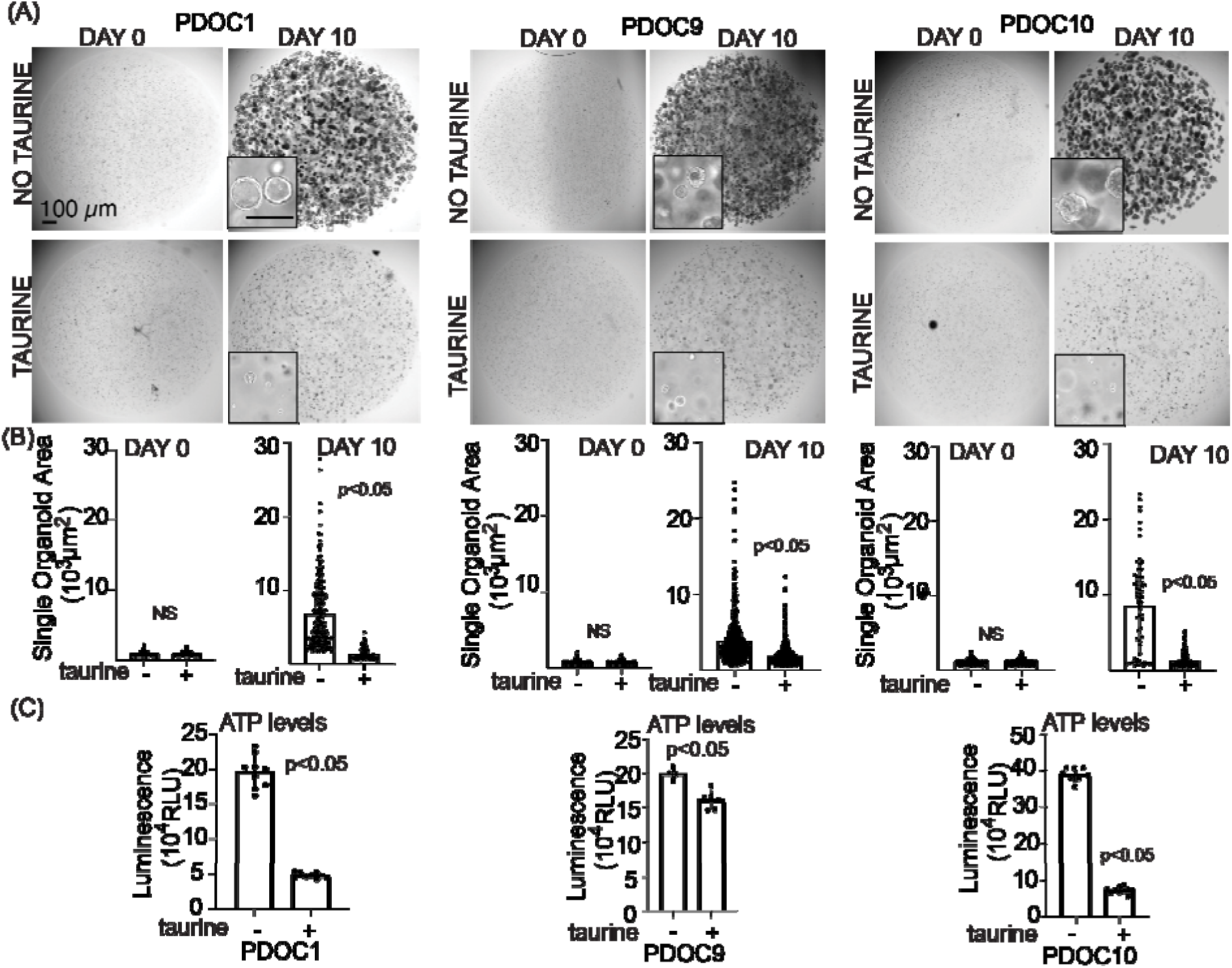
Taurine supplementation suppresses the growth of patient-derived organoids. (A) Low and high (inset) magnification bright-field images of entire wells representing three PDO cultures at Day 0 and Day 10. Inset scale bar is 200 μm. (B) Quantification of single organoid area at day 0 and day 10. 100-700 organoids were quantified per group. Each dot represents one organoid. (C) End-point measurement of ATP levels using CellTiter-Glo® assay. Each dot represents luminescence value recorded from one well. Unpaired, non-parametric two-tailed t- tests with Welch’s correction were used to determine the difference between groups.

### Taurine promotes mutant- and WT- p53 binding to DNA for the activation of p53-mediated cyclin-dependent kinase 1A (p21) expression

To gain mechanistic insight into taurine’s growth and metabolism suppression, we performed transcriptomic analysis of FNE-m-p53 cell monolayer cultures supplemented with taurine and found activation of the p53 pathway (Fig. 5A). These results encouraged us to test whether taurine induces mutant p53 binding to DNA. We used a commercially available p53-DNA binding assay kit (Cayman, #600020) that measures the extent of p53 protein interaction with DNA p53 response elements (Fig. 5B). As a negative control, we used a nuclear extract of CaOv3 cells, which do not express p53, and a positive control was represented by treatment of MCF7 cells (expressing WT p53) with Nutlin (a p53 protein stabilizer). We found that taurine increased the interaction of various mutant p53 proteins with DNA (Fig. 5B).

**Figure 5.**
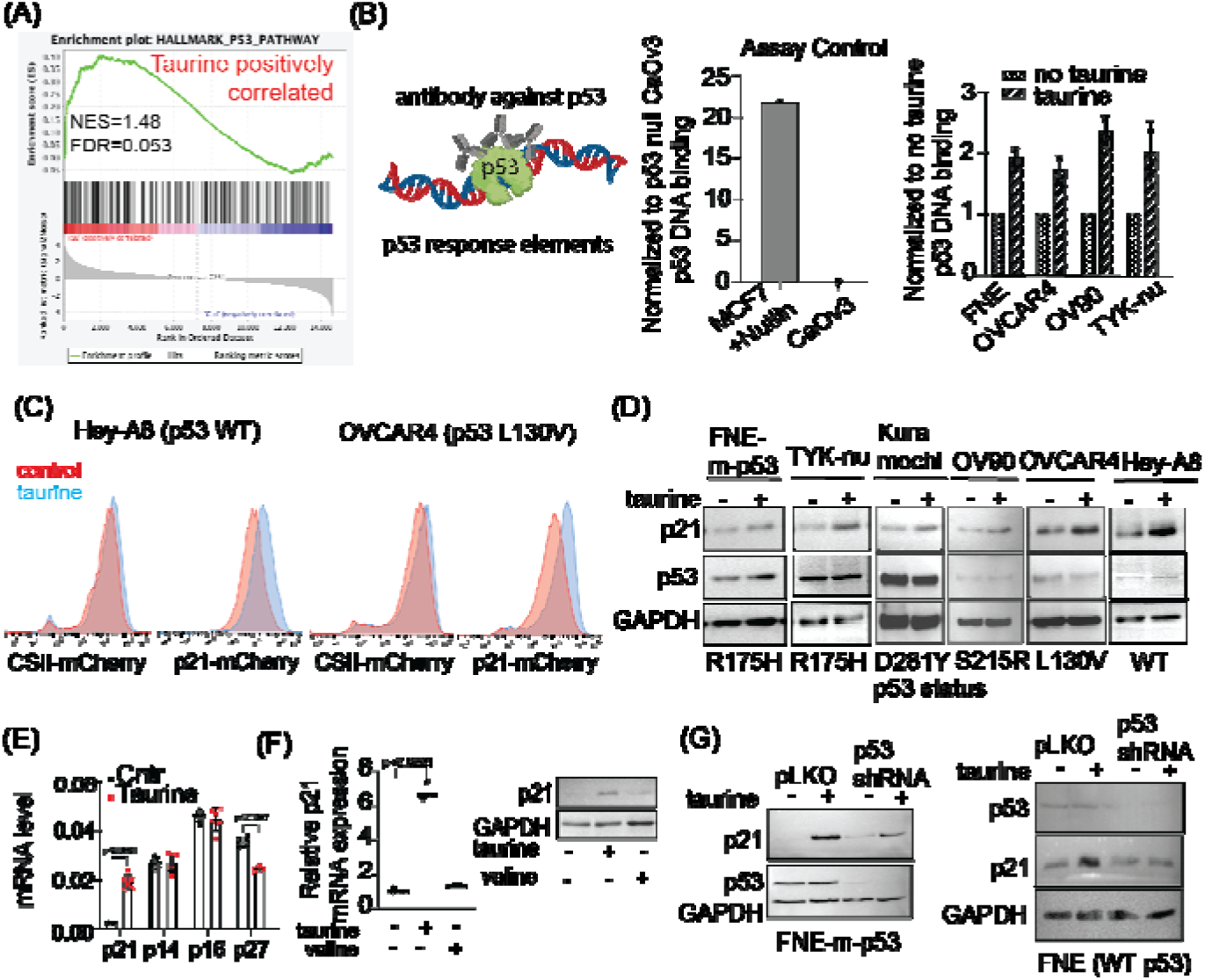
Taurine promotes mutant- and wild-type p53 binding to DNA and p53-mediated p21 activation. (A) Differential transcriptomic analysis of the TP53 gene pathway in FNE-m-p53 cell monolayers grown under normal conditions or in media supplemented with taurine. (B) Quantification of p53 binding to its response element using p53 Transcription Factor Assay Kit (Cayman). For each cell line, data was normalized to the mean of the respective control group and reported as a fold change. (C) Histogram representation of flow cytometry-based analysis of taurine-induced p21 promoter activation. Cells were transduced with a plasmid encoding mCherry cloned downstream of either EF1-α (CSII-mCherry) or the p21 promoter (p21- mCherry). (D) Western blots of p21 and p53 expression in OC cell lines expressing mutant- or wild-type p53. (E) mRNA expression of various cyclin-dependent kinase inhibitors in FNE-m-p53 cells cultured in the absence or presence of taurine. Each dot is one replicate. Data were analyzed using two-way ANOVA. (F) mRNA quantification and western blot analysis of p21 expression in FNE-m-p53 cells cultured in taurine or valine. The p-value represents control *vs.* taurine as determined by one- way ANOVA. (G) Western blots of p21 and p53 expression in FNE-m-p53 and FNE (WT p53) cell lines cultured in taurine following transduction with either scramble shRNA or p53 shRNA.

To extend these studies to living cells, we engineered, based on el-Deiry’s p21 reporter [35], a live p21-mCherry promoter nuclear biosensor, that produces fluorescence in living cells through the engagement of a p53 response element. As a control, we used a CSII-mCherry plasmid, which expresses nuclear mCherry through activity of the EF1-α promoter (Addgene #125262). We found, using flow cytometry, that taurine supplementation to OC cells expressing WT p53 (Hey-A8) or mutant p53 (OVCAR4) increased p21 promoter activity, but not EF1-α promoter activity (Fig. 5C). Consistent with these data, western blot analysis demonstrated taurine-induced p21 protein regulation in a panel of cell lines expressing various mutant p53 proteins or WT p53 protein (Fig. 5D). Taurine supplementation of FNE-m-p53 cell cultures did not induce p14/p16 or p27 transcription (Fig. 5E), and valine failed to stimulate both p21 mRNA and protein expression (Fig. 5F). Importantly, shRNA-mediated depletion of mutant- or WT-p53 expression attenuated taurine-induced p21 activation (Fig. 5G). Furthermore, taurine failed to activate p21 in CaOv3 cells lacking p53 protein expression (S. Fig. 2A). These results support the possibility that p53 (mutant or WT) is partially required for taurine-mediated activation of p21.

In addition to suppression of cell proliferation, we observed that taurine negatively regulated expression of genes supporting glycolysis (Fig. 6A). Western blot analysis revealed activation of the *TP53* inducible glycolysis and apoptosis regulator (TIGAR) (Fig. 6B), a phosphatase that directly inhibits glycolysis [36]. These results suggested a possible inhibition of glycolysis by taurine. To address this possibility, we examined whether taurine inhibits extracellular acidification rate (ECAR) as a proxy of glycolytic activity. ECAR analysis of FNE-m- p53 cell monolayers reveled that taurine suppressed glycolysis and glycolytic capacity, the maximum rate of conversion of glucose to pyruvate/lactate (Fig. 6C and D). Furthermore, we obtained similar results in TYK-nu cell monolayers supplemented with taurine (S. Fig 2B and C). Taken together our results provide evidence that taurine promotes mutant- or WT-p53 interaction with DNA, which is associated with the activation of suppressive mechanisms for growth and metabolism in cultured FNE and OC cells.

**Figure 6.**
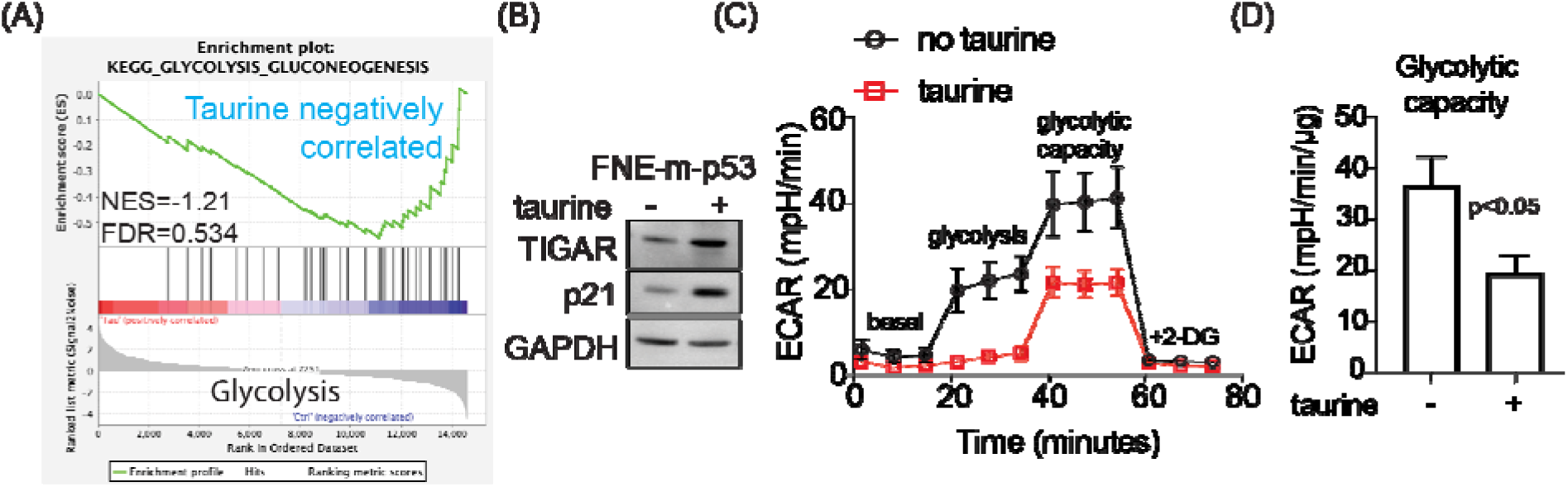
Taurine activates genes involved in the suppression of glycolysis and attenuates glycolytic capacity. (A) Enrichment analysis of transcripts corresponding to genes implicated in the regulation of glycolysis in FNE-m-p53 cells cultured in taurine. (B) Western blot of TIGAR and p21 expression in FNE-m-p53 cells cultured in the absence or presence of taurine. (C) Analysis of ECAR over time showing glycolysis and glycolytic capacity. (D) Bar graph shows average glycolytic capacity with standard deviations (maximal whiskers) across three experiments. An unpaired, two-tailed t-test was used to determine statistical differences between the means.

### Taurine supports cell survival under genotoxic stress

So far, our data are consistent with the idea that taurine supplementation reprograms cells into low-proliferative and reduced-energy metabolic states. Proliferating cells are usually metabolically active and more sensitive to genotoxic stress [37], suggesting a possibility that taurine supplementation, which we demonstrated inhibits proliferation and glycolysis, could protect cells from genotoxic stress. To test this possibility, we used a combination of live cell imaging, flow cytometry, and propidium-iodide- (PI)-based measurements of cell death, taurine supplementation, and DNA damage agent (cisplatin) treatment. We found that taurine supplementation protected FNE-m-p53, FNE cells (WT p53), and additional OC cell lines from genotoxic stress associated with cisplatin treatment (Fig. 7A-B, S. Fig. 3A-B, (MOVIE 2). In accordance with these results, assessment of γH2AX expression (flow cytometry) and DNAPK phosphorylation (western blot) in FNE-m-p53 cells revealed that taurine decreased cisplatin-induced DNA damage (Fig. 7C-E). Furthermore, RNA-seq data analysis of taurine-supplemented FNE-m-p53 cells showed enrichment of genes associated with cell protection from cisplatin, as defined by multiple studies [38–41] (Fig. 7F). Our data support the idea that taurine activates p53 and decreases DNA damage to protect cells from genotoxic stress.

**Figure 7.**
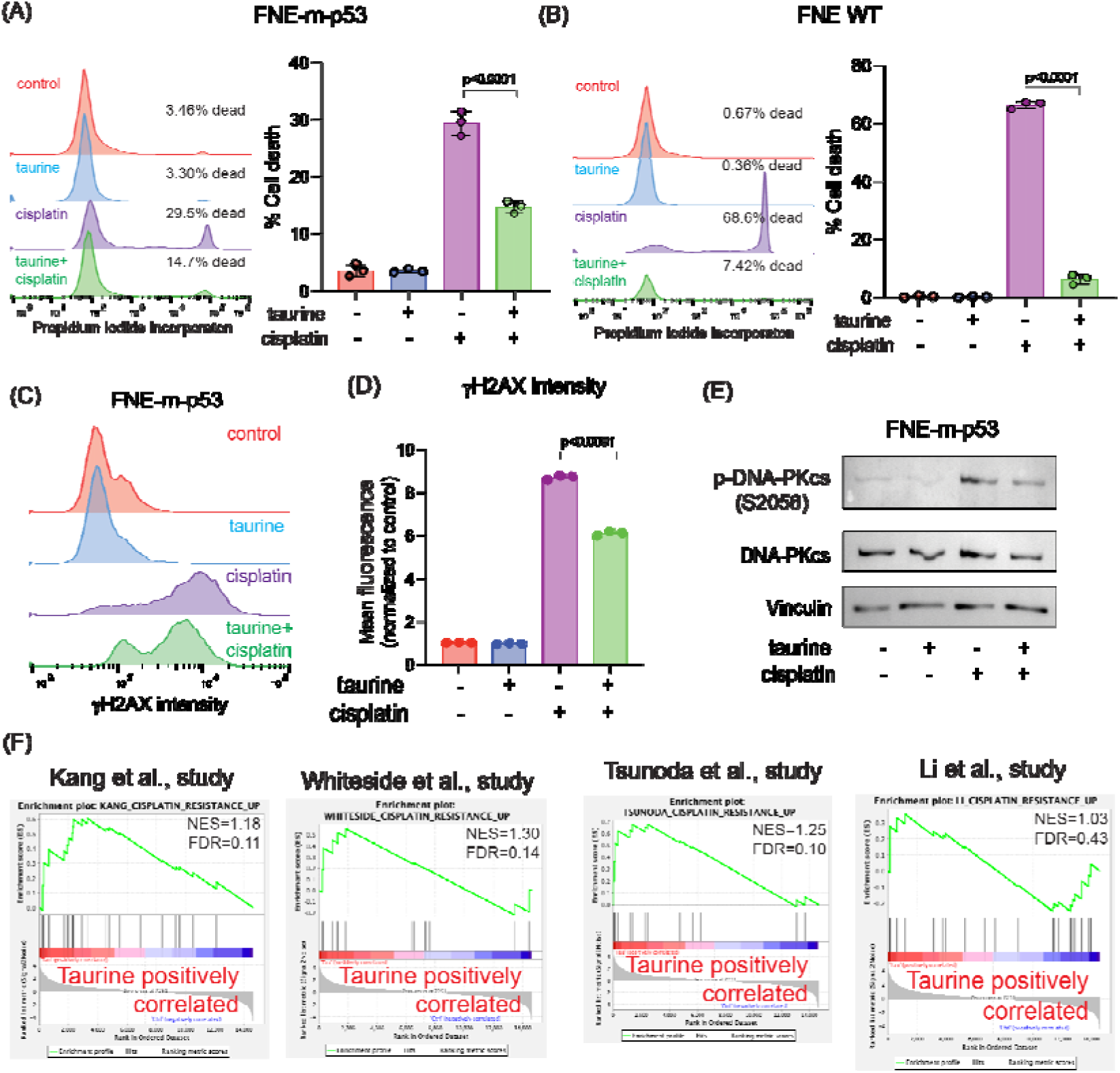
Taurine protects cells both mutant- and wild-type p53 expressing cells from cisplatin-induced cell death and DNA damage. (A-B) Representative histograms and corresponding bar graphs of cell viability for FNE-m-p53 and FNE WT cells cultured in taurine, cisplatin, or combination for 72h. Viability was determined by PI incorporation and quantified using flow cytometry. (C) Representative histograms and (D) quantification of phosphorylated γH2AX staining in FNE-m-p53 cells cultured in taurine, cisplatin, or both for 72h. Each dot is one replicate. Data is normalized to the mean fluorescence of the control group. The p-value differentiates cisplatin *vs*. taurine + cisplatin as determined by one-way ANOVA. (E) Western blot of phosphorylated and total DNA-PKcs protein levels in FNE-m-p53 treated for 72h in the indicated conditions. (F) Enrichment analysis of gene sets associated with cisplatin resistance published in the indicated studies. Taurine treatment of FNE-m-p-53 cells positively correlates with cisplatin resistance.

### Attenuation of p53 does not affect cell protection by taurine

To explore the possibility of p53 involvement in cell protection downstream of taurine, we used p53 shRNA to attenuate the expression levels of the p53 protein. Our experiment revealed that shRNA-mediated attenuation of mutant- or WT-p53 in FNE-m-p53 or FNE cells decreased activation of p21 in response to taurine (Fig. 5G). Next, we wondered whether loss of p53 uncouples taurine from cell protection. Our experiments demonstrated that attenuation of mutant- or WT-p53 expression did not affect taurine s ability to protect cells from cisplatin (Fig. 8A-D). These results suggest that taurine’s cell protection from cisplatin is independent of p53 activation.

**Figure 8.**
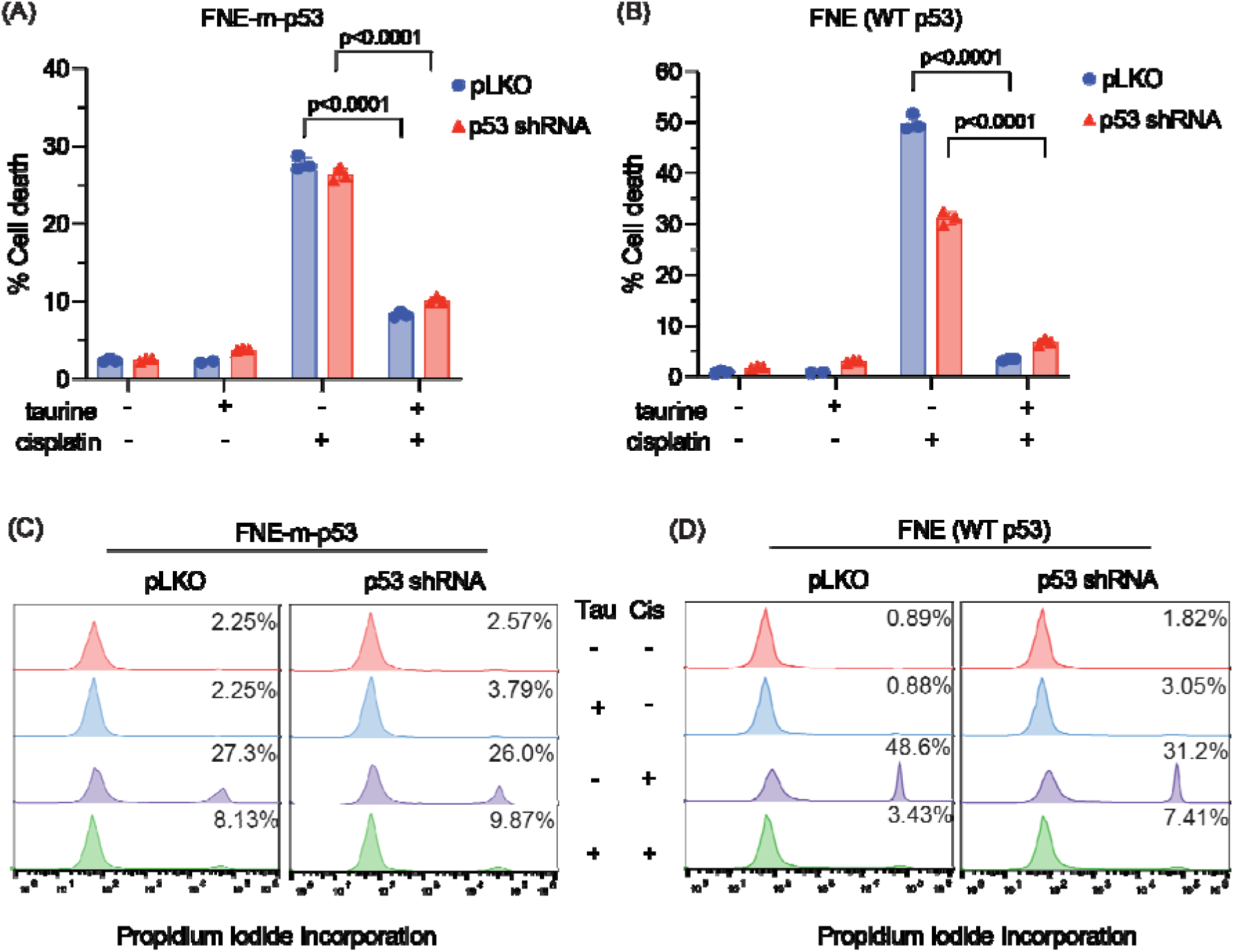
Taurine-induced rescue from cisplatin is not p53 dependent. (A-B) Viability of FNE-m-p53 and FNE (WT p53) cell lines as determined by PI staining and quantified using flow cytometry. Cells were transduced with pLKO scramble shRNA or p53 shRNA and treated with taurine, cisplatin, or both for 72h. Each dot represents one replicate. The p-values represent cisplatin *vs.* taurine + cisplatin for each respective shRNA and were determined by one-way ANOVA. (C-D) Representative histograms from (A) and (B). Percentage of PI positive cells is indicated.

### Taurine mediates cell protection through mTOR/DNAPK/ATM/ATR signal transduction

To identify possible cell protective mechanisms downstream of taurine, we performed reverse phase protein array (RPPA) analysis, to probe activation (expression or phosphorylation) of hundreds of proteins in response to taurine in FNE-m-p53 cells. Our RPPA experiments revealed activation of several major signal transduction pathways, including S6K and ERK (Fig. 9A). We confirmed these results using western blot analysis (Fig. 9B). Furthermore, our RNA-seq analysis indicated that intracellular accumulation of taurine in FNE- m-p53 cells was associated with increased mRNA expression of several DNA damage repair genes, including the Phosphatidylinositol 3-kinase-related kinase (PIKK) family members DNAPK, ATM and ATR (Fig. 9C). Reconciling these results with the activation of S6K, a well-accepted effector of mTOR, which belongs to PIKK family as well, we hypothesized that inhibition of PIKK signaling sensitizes taurine-supplemented cell cultures to cisplatin. To test this, we used Torin2, a clinically relevant ATP-competitive inhibitor of mTOR, DNAPK, ATM/ATR activity [42]. Treatment of FNE-m-p53 monolayers with Torin2 abrogated phosphorylation of S6 ribosomal protein (S. Fig. 3C), and sensitized taurine supplemented cultures to cisplatin (Fig. 10A-B). However, inhibition of mTOR alone with rapamycin did not break taurine protection, nor did inhibition of its downstream effector S6K1 with PF-4708671. Inhibition of ERK with the MEK inhibitor PD03255901 also failed to break taurine’s protection from cisplatin (S. Fig. 3D-H). Our data suggest that taurine protects cells from cisplatin through activation of DNA-damage-sensing mechanisms, including DNAPK and ATM/ATR.

**Figure 9.**
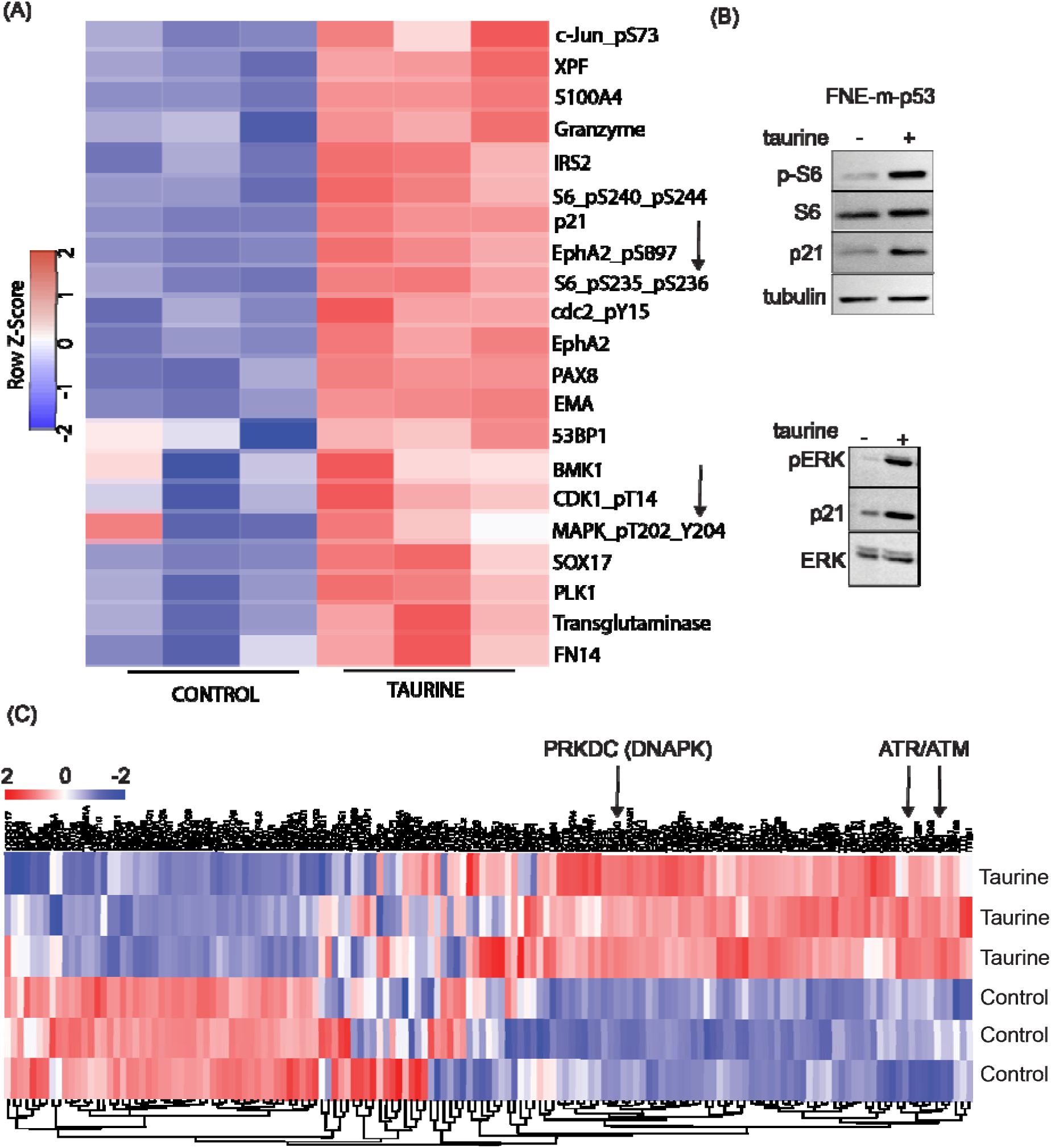
Proteomic and transcriptomic analysis of taurine response. (A) Reverse phase protein array analysis of FNE-m-p53 cells treated with control culture media or media supplemented with 160 mM taurine. Arrows indicate mTOR activity (phosphorylated S6 ribosomal protein) and phosphorylated MAPK/ERK. (B) Western blot for phosphorylated S6 ribosomal protein and phosphorylated ERK for FNE-m-p53 treated with taurine. (C) RNA-seq heatmap for DNA damage response pathways for FNE-m-p53 cells treated with taurine. Arrows indicate PIKK family members PRKDC (DNAPK), ATM, and ATR.

**Figure 10.**
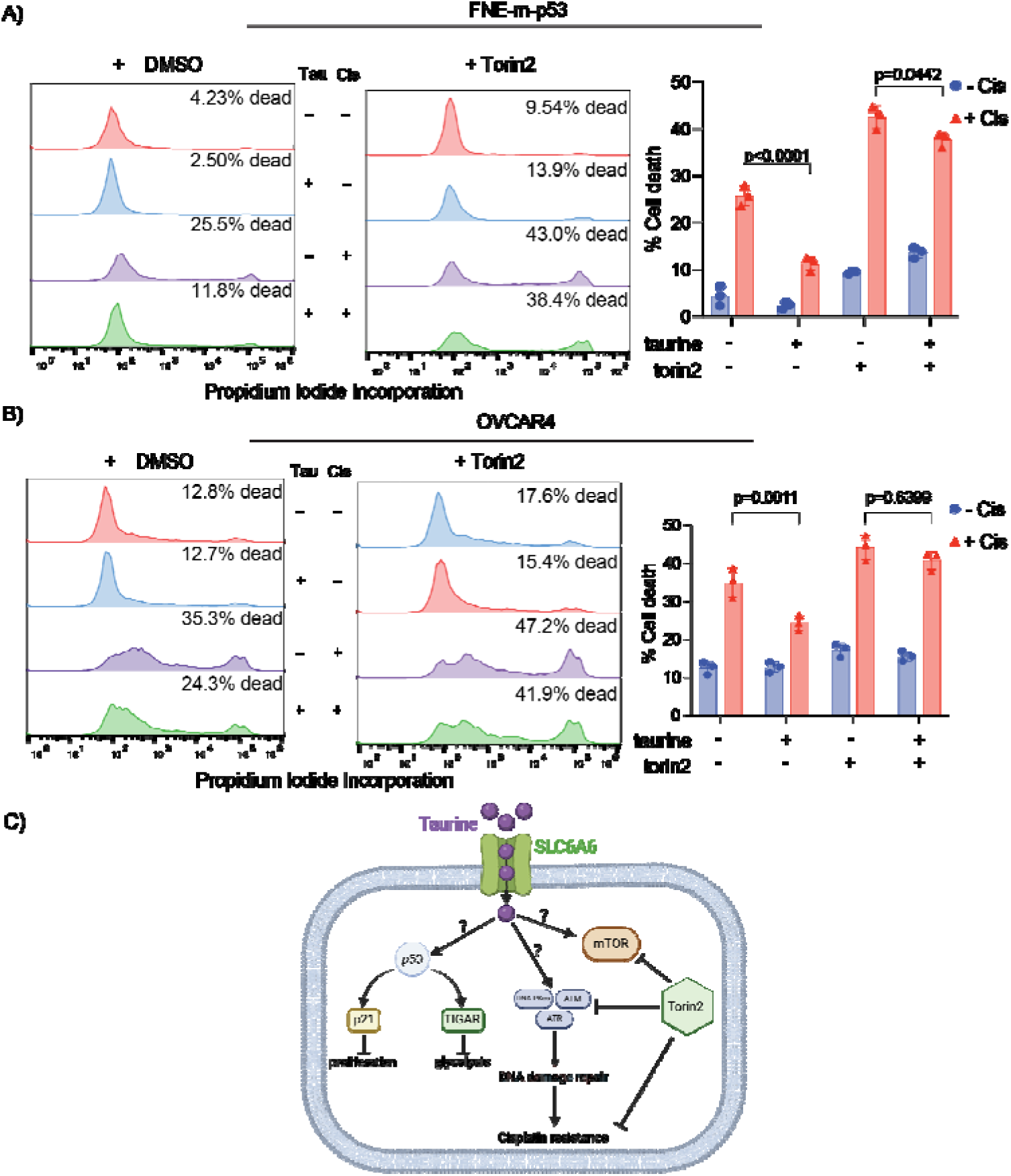
Torin2 sensitizes taurine-treated cells to cisplatin. (A) Representative histograms and quantification of PI incorporation for FNE-m-p53 cells treated under the indicated conditions for 72h. Cells were treated with Torin2 (1μM) for four hours before exposure to cisplatin (10 μM). (B) Representative histograms and quantification of PI incorporation for OVCAR4 cells treated under the indicated conditions for 24h. Data are reported as mean ± SD. p-values represent DMSO + cisplatin vs. DMSO + taurine + cisplatin and Torin2 + cisplatin *vs.* Torin2 + taurine + cisplatin. Statistical analysis was performed using two-way ANOVA. (C) Schematic representing SLC6A6- mediated intracellular taurine accumulation and activation of cell signaling pathways involved in suppression of proliferation and glycolysis, and response to DNA damage.

## DISCUSSION

These studies provide potentially new insights into the mechanisms of cell growth suppression, and cell protection by taurine, a non-proteogenic sulfonic amino acid present in the cytoplasm of ascites cells derived from OC patients. Mass-spectrometry-based quantitative examination of intracellular taurine pools revealed taurine concentrations ranging from 45 μM to 900 μM among ascites cells. These taurine levels were on average 100-fold higher than taurine intracellular pools found in cultured cells. Mimicking ascites cell-associated intracellular taurine concentrations in cell culture suppressed cell proliferation, glycolysis, and induced cell protection from genotoxic stress. Taurine decreased cisplatin-mediated DNA damage, indicating that intracellular accumulation of the free amino acid taurine might contribute to the evolution of slowly proliferating cell clones with increased ability to repair DNA upon stress. Combination of bulk RNA-seq, and proteomic (RPPA) analysis of taurine-supplemented FNE-m-p53 cell cultures revealed activation of the p53, ERK and mTOR pathways, and increased mRNA expression of DNAPK/ATM/ATR. Taurine induced mutant- or wild-type p53 binding to DNA, and the activation of p53 effectors involved in the negative regulation of the cell cycle and glycolysis, including p21 and TIGAR, respectively. Hairpin RNA-mediated attenuation of p53 decreased activation of p21, but it did not affect taurine’s ability to protect cells from cisplatin. Similarly, inhibition of ERK pathway by MEK inhibitor PD03255901 did not alter taurine’s protective activities. However, treatment of taurine supplemented cell cultures with Torin2, a clinically relevant ATP-competitive inhibitor of mTOR, DNAPK and ATM/ATR was sufficient to block taurine-mediated cell protection. Our data are consistent with a model whereby the accumulation of intracellular taurine suppresses proliferation and provides cell protection from genotoxic stress through signal transduction pathways involving mTOR/DNAPK/ATM/ATR.

Proliferative ovarian tumors, as assessed by Ki67 stains, are initially sensitive to chemotherapy treatments [43]. However, the disease recurs, and its progression has been recently linked to the evolution of slowly proliferating tumor cell clones [44], indicating that cell protection from treatment could be linked to molecular mechanisms associated with suppression of proliferation. A set of new studies provide evidence that activation of p53-mediated suppression of the cell-cycle and metabolism can be associated with cell survival under treatment [15, 45, 46]. However, the role of intracellular taurine in p53 activation and therapy resistance is not completely understood, nor is it clear whether the activation of p53 in response to taurine is also associated with induction of pro-survival mechanisms. Here we provide evidence that taurine suppresses OC cell proliferation and activates a p53 effector, p21, a molecule directly involved in cell-cycle arrest. However, p53 activation was not required for taurine-induced cell protection from cisplatin, indicating that p53 is dispensable in taurine cell protection.

Taurine is an osmolyte [47] and recent results support the idea that accumulation of intracellular osmolytes promotes cell-size increase and cell-cycle arrest [34]. Our cell culture data demonstrate that elevation of intracellular taurine to the levels found in cells isolated from OC ascites significantly increases OC cell size, and this phenotype is associated with suppression of proliferation. Based on these results, it is possible that one of the mechanisms supporting evolution of growth-suppressed tumor cell clones involves influx of free amino acid osmolytes, and cell size changes.

Recent *in silico* and cell-free biochemical studies [48] demonstrated that taurine can directly bind cyclin-dependent kinase 6 (CDK6). This interaction appears to inhibit CDK6 activity in solution, supporting the possibility that intracellular taurine accumulation could directly suppress CDK6/Cyclin D activity, which could lead to activation of p53 [49, 50] and inhibition of proliferation. Our experiments also provide evidence that taurine increases mutant- or wild-type p53 binding to DNA. Biochemical evidence exists [51] that taurine interacts with casein proteins to change their conformation. Thus, it is possible that taurine could directly interact with p53 and change its conformation to support DNA binding and subsequent activation of effector molecules including p21 and TIGAR. However, taurine’s ability to protect cells from cisplatin did not depend on p53 but required mTOR/DNAPK/ATM/ATR activity. These results suggested the idea that emergence of therapy-resistant cell clones is driven by the mechanisms of cell-cycle suppression and the activation of major cell survival and DNA-repair signal transduction pathways including mTOR/DNAPK/ATM/ATR. How taurine activates mTOR/DNAPK/ATM/ATR is not known, and our data does not support that taurine itself induces DNA damage (as measured by γH2AX levels), implicating existence of taurine-induced DNA-damage-independent mechanisms of transcriptional regulation of DNA-damage-sensing kinases.

In conclusion, our studies provide a possible mechanism whereby intracellular accumulation of taurine can activate p53 and mTOR/DNAPK/ATM/ATR signaling pathways to suppress cell growth and support the emergence of slowly proliferating cells resistant to DNA-damage-based therapies (Fig 10C). We identified that taurine-mediated cisplatin protection was abolished by treatment with Torin2, a clinically relevant small molecule inhibitor of mTOR/DNAPK/ATM/ATR.

## Supplementary Materials

S. Fig 1: Analysis of proliferation and cell-cycle in OC cell lines. S. Fig 2: Western blot analysis of p21 expression in CaOv3 cells and ECAR analysis of TYK-nu cells. S. Fig 3: Propidium iodide incorporation of Hey-A8 and OVCAR4 cells. Western blot and propidium iodide incorporation for FNE-m-p53 treated with inhibitors for MEK, mTOR, and S6K1. Movie 1: Time lapse of FNE-m-p53 treated with control media or 160 mM taurine for 72h. Movie 2: Time lapse of FNE-m-p53 cells treated with cisplatin (10 μM) or taurine and cisplatin for 72h.

## Author contributions

Daniel Centeno: Conceptualization, Methodology, Validation, Formal analysis, Investigation, Writing - original draft, Visualization. Sadaf Farsinejad: Conceptualization, Methodology, Software, Formal analysis, Data curation, Visualization. Elena Kochetkova: Conceptualization, Methodology, Formal analysis, Investigation, Funding acquisition. Tatiana Volpari: Methodology, Investigation, Formal analysis. Aleksandra Gladych- Macioszek: Methodology, Investigation, Formal analysis. Agnieszka Klupczynska – Gabryszak: Methodology, Investigation, Formal analysis. Teagan Polotaye: Validation, Formal analysis, Investigation. Michael Greenberg: Validation, Investigation. Douglas Kung: Conceptualization, Validation, Investigation. Emily Hyde: Validation, Investigation. Sarah Alshehri: Conceptualization. Tonja Pavlovic: Concenptualization. William Sullivan: Resources. Szymon Plewa: Resources. Helin Vakifahmetoglu-Norberg: Resources. Frederick J Monsma, Jr: Resources. Patricia A. J. Muller: Resources, Writing - review & editing. Jan Matysiak: Resources. Mikolaj Zaborowski: Resources. Analisa DiFeo: Resources. Laura A. Martin: Conceptualization, Methodology, Funding acquisition. Erik Norberg: Conceptualization, Methodology, Funding acquisition. Marcin Iwanicki: Conceptualization, Methodology, Writing - original draft, Visualization, Supervision, Funding acquisition.

## Funding

This study was supported by NIH/NCI R21CA256615 (M.I.), an Olipass Corporation grant (M.I.), the Kaleidoscope of Hope Ovarian Cancer Research Foundation (M.I.)., the New York Stem Cell Foundation Research Institute (NYSCF), the NCI R21CA240219 (L.A.M.). The Swedish Research Council 2021-01787, The Swedish Cancer Society CAN 2017/466, CAN 2017/1015, 20 0979 PjF (E.N.), Stiftelsen Lars Hiertas Minne FO2021-0082, Stiftelsen Längmanska kulturfonden BA22-094, Stiftelsen Sigurd & Elsa Goljes Minne LA2022-0075 (E.K.). Tissue samples were provided by the NCI Cooperative Human Tissue Network (CHTN) RRID SCR_004446. Other investigators may have received specimens from the same tissue specimens.

## Institutional Review Board Statement

All patients provided a signed informed consent, approved by the Ethics Review Board of Poznań University of Medical Sciences (Consent No 737/17). All procedures performed in studies involving human participants were in accordance with the ethical standards of the institutional and national research committee and with the 1964 Helsinki declaration and its later amendments or comparable ethical standards.

## Informed consent statement

Informed consent was obtained from all subjects involved in the study.

## Acknowledgments

We would like to thank Thomas Cattabiani for editorial help with the manuscript preparation. D.C. is supported by the Ajay Bose Memorial Scholarship and Robert Crooks Stanley Graduate Fellowship.

## Conflict of Interest

The authors declare no conflict of interest.

## Supplementary Materials

**Supplemental Figure 1.**
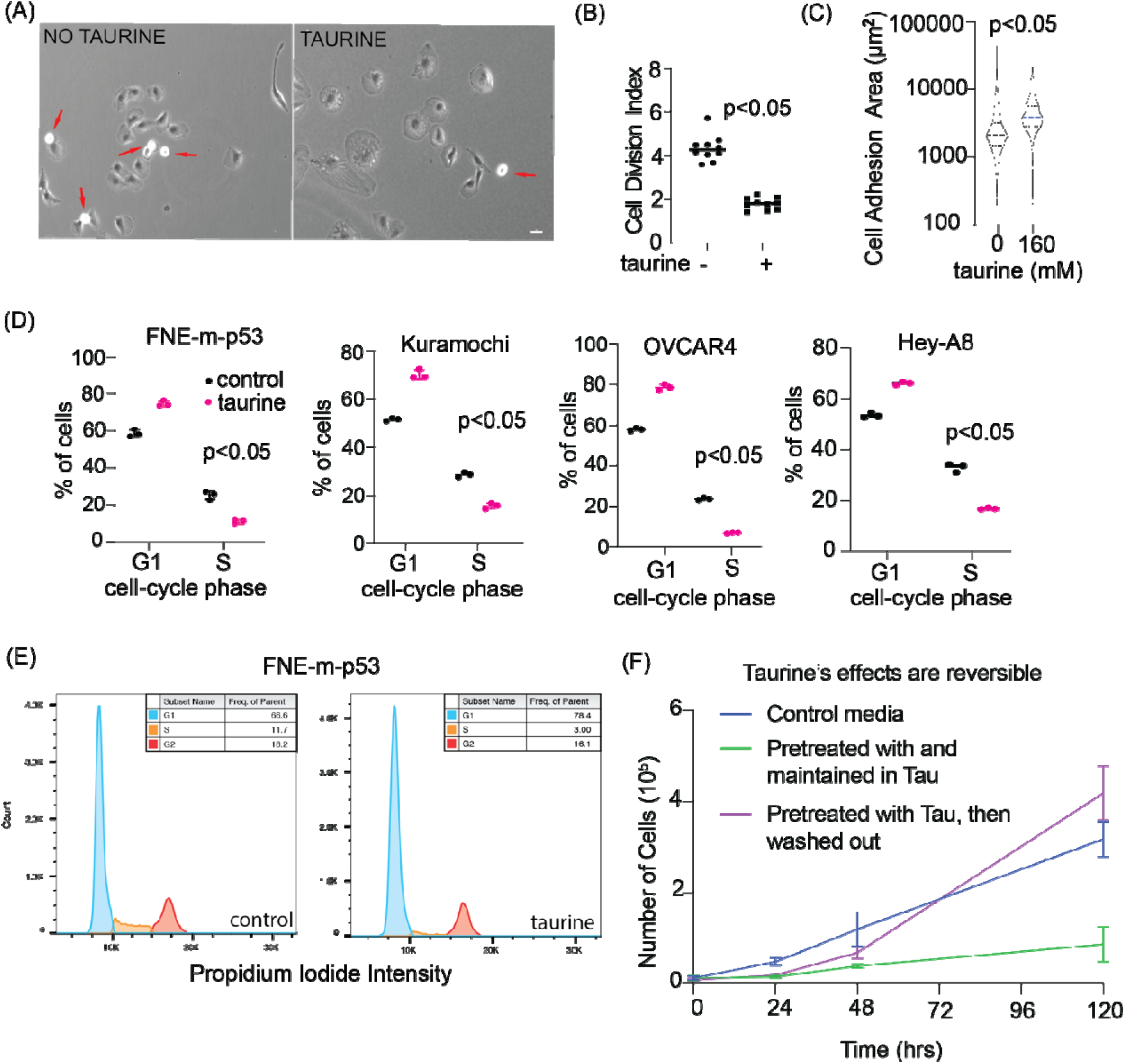
(A) Phase-contrast images of FNE-m-p53 cells. The red arrows point to mitotic cells. Scale bar is 50 µm. (B) Quantification of FNE-m-p53 cell division based on live-cell fluorescent imaging. Cells were tracked for 72h using TrackMate. Cell division index is calculated as the number of tracks (cells) detected over time divided by the number of track splits (cell divisions). Each dot represents the mean index of one ROI. Statistical analysis was performed using student’s t-test. (C) Quantification of GFP-expressing FNE-m-p53 cell adhesion area based on GFP signal obtained from live-cell microcopy. Far the control group, n=2994 cells, and for taurine treatment, n=1164 rails. Statistical analysis was performed using student’s t-test with Welch’s correction. (D) Flow cytometry and Pl-based cell-cycle analysis of cell lines treated with 160 mM taurine for 72h. Cell Cycle function In FlowJo was used to determine the frequency each cell-cycle phase. Each dot is one replicate. Bars indicate SD. Student’s t-test was used to determine statistical significance. (E) Representative histograms for FNE-m-p53 cells shown in (D). (F) Quantification of cell proliferation based on automated counting. Cells were treated with control media or media with taurine for three days. Cells were then replated, and taurine treated cells were either washed and placed in control media or maintained in taurine for the indicated time. Cells were counted in triplicate.

**Supplemental Figure 2.**
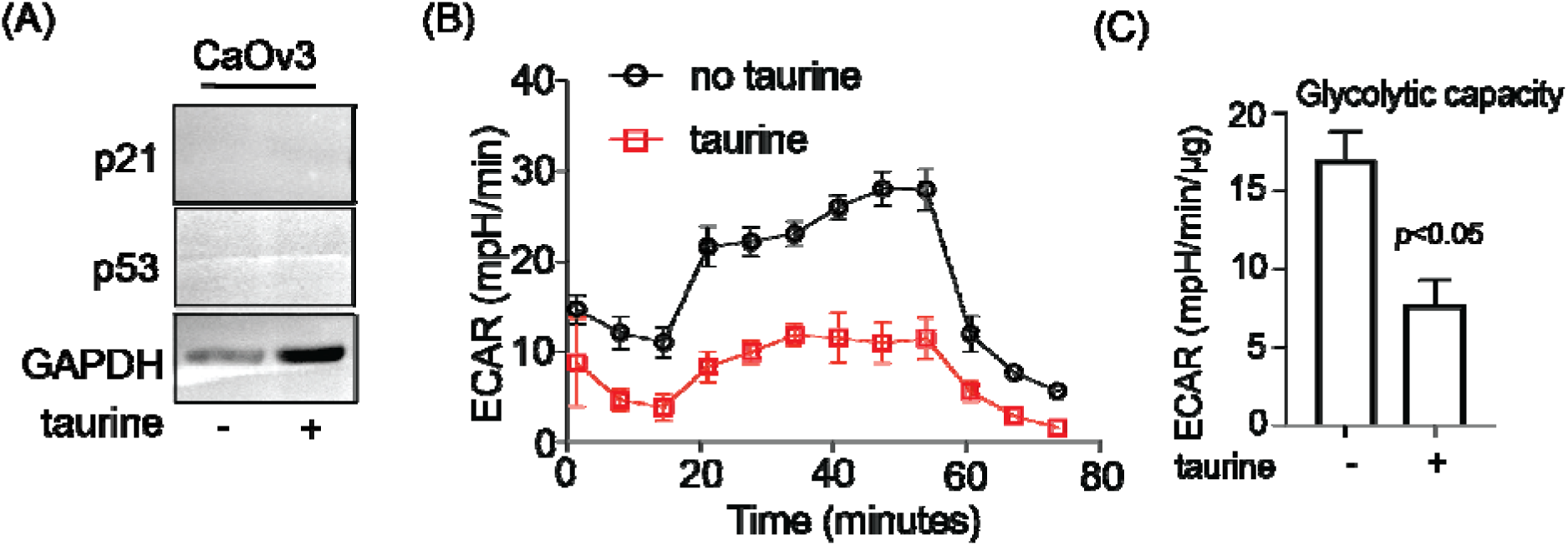
(A) Western blot for p21 and p53 in CaOv3 cell monolayers treated with 160 mM taurine for 72h. (B) Analysis of ECAR over time representing giycoiysis and (C) glycolytic capacity in TYK-nu cell monolayers treated with 160mM taurine. Bar graph shows the average values with standard deviations (maximal whiskers) across three experiments. An unpaired, two-tailed t-test was used to determine statistical differences between the means.

**Supplemental Figure 3.**
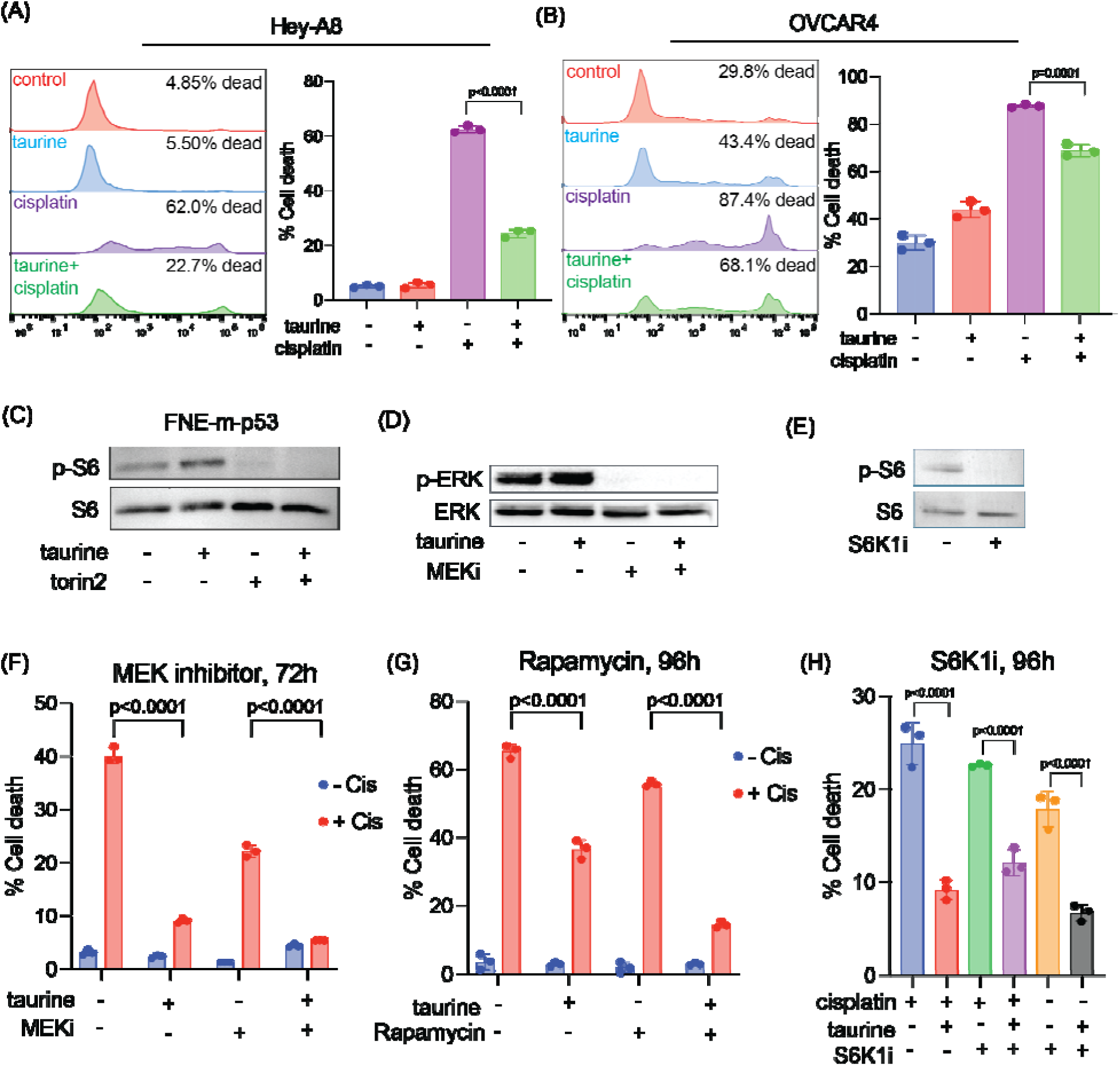
(A-B) Representative histograms and quantification of PI incorporation of Hey-AB (A) and OVCAR4 (B) cells treated under the indicated conditions for 72h. Statistical significance was determined by two-way ANOVA p-values represent cisplatin vs. taurine + cisplatin treated groups. Each dot is one replicate. (C-E) Western blot of FNE-m-p53 treated with Torin2 (C), MEK inhibitor (D) or S6K1 inhibitor (E). (F-H) Quantification of PI incorporation in FNE-m-p53 cells treated with a MEK inhibitor (F), rapamycin (6), or S6K1 inhibitor under the indicated conditions. Each dot is one repicate. Statistical significance was determined by ANOVA For all bar graphs, data are presented as mean ± SD.

## Notes

### Competing Interest Statement

The authors have declared no competing interest.

### Summary of Updates

The manuscript has been extensively revised, and includes additional data that support taurine's tumor suppressive effects. Furthermore, the manuscript has been updated and provides mechanistic insights on taurine-induceded protection from chemotherapy. Last, a mislabeling of a cell line was corrected. All figures were revised.

